# The NS1 protein of contemporary West African Zika virus is efficient to increase cellular permissiveness to virus replication

**DOI:** 10.1101/2024.04.10.588981

**Authors:** Machmouchi Dana, Courageot Marie-Pierre, Ogire Eva, Kohl Alain, Philippe Desprès, Roche Marjolaine

## Abstract

Mosquito-borne Zika virus (ZIKV; orthoflavivirus, *Flaviviridae*) has become a global health problem due to expansion of the geographic distribution of Asian Lineage virus. Contemporary ZIKV strains of African lineage have recently gained increased attention due to their epidemic potential and their capacity to be highly teratogenic in humans. The ZIKV non-structural NS1 protein from recent West African strains Africa was been studied where with view of its importance in the pathogenicity. NS1 protein from contemporary West African ZIKV (NS1^CWA^) and historical African ZIKV strain MR766 (NS1^MR766^) differ by seven amino-acid substitutions. Expression of recombinant NS1 proteins showed differences in the subcellular distribution between NS1^CWA^ and NS1^MR766^ in HEK-293T cells. There was an increased secretion efficiency of soluble NS1^CWA^ compared to NS1^MR766^. The replication of a chimeric MR766/NS1^CWA^ virus was studied in Vero and A549 cells. Insertion of NS1^CWA^ into MR766 enhances virus replication in both cell lines leading to more pronounced cell death. This correlated with lower up-regulation of *IFN-β* and interferon-stimulated gene mRNA in A549 cells infected by MR766/NS1^CWA^ virus. Our data raise the question on the importance of NS1 protein in the pathogenicity of contemporary ZIKV from West Africa, and point to differences within viral strains belonging to the same African lineage.

**AUTHOR SUMMARY:** Mosquito-borne Zika virus (ZIKV) of African lineage has the potential to cause epidemic along with a high risk of fetal pathogenicity. Too little is still known on the features of contemporary ZIKV from West Africa. We find there is a remarkable conservation of NS1 amino-acid residues between ZIKV strains recently isolated in Senegal and Guinea. Analysis of recombinant ZIKV NS1 protein revealed efficient secretion of contemporary African NS1 protein from human cells. Using infectious molecular clone of African ZIKV, we showed that contemporary West Africa NS1 protein influences virus replication and innate immune activation. The NS1 protein has been proposed as playing a major role in the pathogenicity of contemporary ZIKV from West Africa.

## INTRODUCTION

The mosquito transmitted Zika virus (ZIKV) belongs to orthoflavivirus genus of *Flaviviridae* family has become an increasingly important global health problem (1). In the past decade, the expansion of the geographic distribution of ZIKV of Asian genotype and rapid spread caused major epidemics in the South Pacific in 2013, and for the first time in South America in 2015 (1,2). Emergence of Asian ZIKV strains in South Pacific and then Western Hemisphere has been associated with Guillain-Barré syndrome and unprecedented teratogenic effects presenting as microcephaly and other developmental and neurological disorders (1,3-5). Although ZIKV transmission in adults classically involves a blood meal by infected female mosquitoes of the *Aedes* species, sexual contact, blood transfusion and intrauterine transmission have been documented as non-vectored transmission routes (1).

Studies aiming to understand the features of emerging ZIKV were mostly carried out using epidemic strains of Asian lineage that had been firstly isolated in 2013 (South Pacific) and then 2015-16 (South America and Caribbean islands). Although African ZIKV strains should also have epidemic potential, and high risk of fetal pathogenicity has been demonstrated (6-9), there is little information on molecular characteristics of contemporary ZIKV of African lineage. Among ZIKV non-structural (NS) proteins, NS1 glycoprotein (352 amino-acid residues) exists as a membrane-associated homodimer in the endoplasmic reticulum (ER) where the protein associates with other NS proteins into viral replication complexes (10-12). A part of a hydrophobic NS1 dimer is driven towards the cell membrane where the protein is present at the cell surface whereas it can also be released as lipoprotein particle playing a role in the pathophysiology of ZIKV infection (13-15). The NS1 protein plays a major role in evading host innate immunity and also influences virus acquisition by mosquitoes during a blood meal (16-20).

Contemporary West African ZIKV strain ZIKV-15555 has been sequenced from an individual who has been infected in Guinea in 2018 whereas viral strains Senegal-Kedougou 2011, and Senegal-Kedougou 2015 have been isolated from mosquito pools in Senegal in 2011 and 15, respectively (6). Analysis of their RNA genomic sequences showed full amino-acid conservation in the NS1 protein. Here, we characterized contemporary West Africa ZIKV NS1 protein (intitled hereafter NS1^CWA^) based on a comparative study with NS1 from the historical viral strain MR766 (intitled hereafter NS1^MR766^), isolated from a non-human primate in Uganda in 1947. Expression of NS1^CWA^ and NS1^MR766^ proteins which differ by seven amino-acid substitutions was examined using recombinant NS1 proteins. Analysis of recombinant ZIKV NS1 protein revealed efficient secretion of NS1^CWA^ from human cells. Using a chimeric MR766 virus bearing NS1^CWA^ sequence, we showed that NS1^CWA^ is efficient to enhance virus replication and down-regulate innate immune activation.

## METHODS

### Cells and antibodies

Human embryonic kidney HEK-293T (ATCC, CRL-1573), human carcinoma epithelial lung A549 (Invivogen Inc, Toulouse, France), and Monkey kidney normal Vero E6 (CCL-81, ATCC, Manassas, VA, USA) cells were grown in Dulbecco’s modified Eagle’s medium (DMEM) growth medium supplemented with heat-inactivated fetal calf serum (FBS, Dutscher, Brumath, France), respectively, and antibiotics (PAN Biotech Dutscher, Brumath, France) at 37 °C in a 5% CO_2_ atmosphere. The purified mouse anti-*pan* flavivirus envelope E protein monoclonal antibody (mAb) 4G2 was produced by RD Biotech (Besançon, France). The purified mouse anti-NS1 mAb 4G4 was a generous gift from Dr D. Watterson (Brisbane, Australia). Rabbit anti-FLAG antibody obtained from DIAGNOMICS (Blagnac, France) was used to detect recombinant FLAG-tagged protein. Immunoblot assay on ISG was performed using anti-ISG15 and anti-IFIT1 antibody (Thermo Fisher Scientific Illkirch-Graffenstaden, France). Donkey IgG anti-rabbit IgG-Alexa Fluor 488 and anti-mouse IgG-Alexa Fluor 594 were used as secondary antibodies for immunofluorescence analysis.

### Construction of a chimeric MR766 virus

The infectious molecular clone MR766^MC^ that has been obtained from African ZIKV strain MR766-NIID (GenBank accession number LC002520) was previously described by Gadea *et al*. (2016) (21). Based on the genetic reverse method ISA, production of MR766^MC^ involves three synthetic genes Z-1^MR766^, Z-23^MR76^ and Z-4^MR766^ cloned into pUC57 plasmid. The gene Z-1^MR766^ includes the CMV promoter immediately adjacent to the 5’NCR followed by the coding region for the structural proteins. The Z-23^MR766^ gene encodes the nonstructural proteins NS1 to NS4B. The gene Z-4^MR766^ encodes the NS5 protein followed by the 3’NCR and ended by hepatitis delta virus ribozyme and SV40 poly(A) signal. The three fragments were amplified by PCR from their respective plasmids using a set of specific primers so that Z-1^MR766^ and Z-23^MR766^ as well as Z-23^MR766^ and Z-4^MR766^ matched on at least 30 nucleotides. To generate MR766^MC^, the three purified PCR products are transfected into HEK-293T cells using Lipofectamine 3000 and after 4 days, cell supernatant was recovered and used to infected Vero E6 cells in a first round of amplification (P1). After 5 days, P1 was recovered and amplified for a further 3 days to produce a working virus stock P2. Virus stock titers are determined by a standard plaque-forming assay as previously described (21,22,24). Infectious virus titers are expressed as plaque-forming units per ml (PFU.mL^-1^). To generate a chimeric MR766^MC^ virus bearing the gene coding for ZIKV-15555 NS1 protein, a Z-23^ZIKV-15555^ gene coding for ZIKV-15555 NS1 to NS4B proteins (GenBank accession number MN025403) was synthetized and inserted into plasmid pUC57 by Genecust (Boynes, France). The fragment of Z-23^ZIKV-15555^ gene coding for NS1 protein was amplified by PCR using a couple of specific primers (Table S1). Site-directed mutagenesis by PCR was conducted on Z-23^MR766^ gene to generate a sub-fragment Z-23^MR766(NS2A/4B)^ gene coding for NS2A to NS4B proteins. HEK-293T cells were transfected with four PCR products amplified from Z-1^MR766^, NS1^ZIKV-15555^, Z-23^MR766(NS2A/4B)^, and Z-4^MR766^ genes. The extremities of NS1^ZIKV-15555^ can match with the 3’end of Z-1^MR766^ and the 5’end of Z-23^MR766(NS2A/4B)^ preserving the large open reading frame of viral polyprotein. The recovered chimeric MR766-(NS1^ZIKV-15555^) virus was twice amplified on Vero cells as described above and a working virus stock P2 was used for the further study.

### Recombinant ZIKV NS1 proteins

Mammalian codon-optimized genes coding for the transmembrane domain 2 of ZIKV E protein acting as authentic NS1 signal peptide followed by the residues 1/352 of the ZIKV NS1 strains MR766 (GenBank accession number LC002520) and ZIKV-15555 (GenBank accession number MN025403) were established using *Homo sapiens* codon usage as reference. A glycine-serine spacer followed by a FLAG tag were inserted in-frame at the C-terminus of recombinant NS1 protein. The synthesis of gene sequences and their cloning into *Nhe*-I and *Not*-I restriction sites of the pcDNA3.1-hygro (+) vector plasmid to generate recombinant plasmids pcDNA3/NS1^MR766^ and pcDNA3/NS1^cWA^ were performed by Genecust (Boynes, France). Site-directed mutagenesis was conducted on pcDNA3/NS1^MR766^ to introduce the two NS1 amino-acid substitutions S92P and Y286H. The resulting plasmid pcDNA3/NS1^MR766^-(P92, H286) was obtained by Genecust (Boynes, France). The plasmid sequences were verified by Sanger method. Endotoxin-free plasmid DNA purification was performed by Genecust (Boynes, France). HEK-293 T cells were transient transfected with plasmids using Lipofectamine 3000.

### RT-qPCR

Total RNA was extracted from cells using RNeasy kit (Quiagen) and reverse transcription was performed using random hexamer primers (intracellular viral RNA) and MMLV reverse transcriptase (Life Technologies). Quantitative PCR was performed on a ABI7500 Real-Time PCR System (Applied Biosystems, Life Technologies, Villebon-sur-Yvette, France). Data were normalized using 36B4 as RPLP0 housekeeping gene. For each single-well amplification reaction, a threshold cycle (*C*t) was calculated using the ABI7500 program (Applied Biosystems, Life Technologies) in the exponential phase of amplification. Relative change in gene expression was determined using the 2∂∂*C*t method and reported relative to the control. The couples of primers for amplifying house-keeping 36B4, ISG and IFN-β mRNA and ZIKV genomic RNA (E gene) are listed in Table S1.

### Immunoblot assay

Cell lysates were performed in RIPA lysis buffer (Sigma, Lyon, France). Proteins were separated by 4–12% SDS–PAGE and transferred into a nitrocellulose membrane. After blocking of the membrane for 1 h with 90% FBS or 5% milk in TBS-Tween, blots were incubated with primary antibody at dilution 1:200. Anti-mouse or anti-rabbit IgG HRP-conjugated secondary antibodies were used at 1:5000 dilution. For dot blot assays, samples were directly loaded on a nitrocellulose membrane and then probed with the primary antibody and then anti-mouse or anti-rabbit IgG HRP-conjugated secondary antibody. Membranes were developed with Pierce ECL Western blotting substrate (Thermo Fisher Scientific, Les Ulis, France) and exposed on an Amersham imager 680 (GE Healthcare). The signal intensity of NS1 was measured by Image J software. The results are the mean of two or three independent assays, as indicated. The protein abundance ratio between NS1 and β-actin was determined for monomeric and dimeric forms.

### Confocal immunofluorescence assay

HEK-293T cells seeded on coverslips were fixed with 3.7% paraformaldehyde (PFA) in PBS. Permeabilization of fixed cells was performed for 4 min with nonionic detergent Triton X-100 at the final concentration of 0.1% in PBS. Cells were stained with primary FLAG antibody at dilution 1:1000 in PBS containing 1% bovine serum albumin (BSA) for 1 h at RT. The goat anti-rabbit Alexa Fluor 488 IgG was used as the secondary antibody (1/1000) for 30 min in the dark. After washing, nucleus morphology was revealed by DAPI staining (final concentration 100 ng/mL). Image acquisition was carried out with a Zeiss LSM 710 laser scanning confocal microscope (LSM 710, Zeiss, Germany) equipped with a X63 objective. Each *z*-stack was then processed using Amira® 6.1 (FEI, Mérignac, France) to obtain a surface rendering image for one or two fluorescent markers.

### Flow cytometry assay

For flow cytometry analysis, cells were harvested after trypsinization and fixed with 3.7% PFA in PBS at RT for 10 min. A solution of Triton X-100 (0.15%) in PBS was used to permeabilize the fixed cells for 5 min at RT. After incubation of cells with a blocking solution for 10 min, ZIKV infectivity was assessed using the mouse anti-E protein mAb 4G2 (RD-Biotech, Besançon, France). Antibody donkey anti-mouse Alexa Fluor 488 IgG (Invitrogen) at dilution 1:2000 served as secondary antibody. Immunostained cells were subjected to flow cytometric analysis using FACScan flow cytometer (CytoFLEX, Beckman Coulter, Brea, CA, USA). For each assay, at least 10,000 cells were analyzed and the percentage of positive cells was determined using CytExpert software (version 2.1.0.92, Beckman Coulter, Brea, CA, USA).

### Cytotoxic assay

For lactate dehydrogenase (LDH) assay, cells were seeded in 12-well culture plates. Cytotoxicity was evaluated by quantification of lactate dehydrogenase (LDH) release in cell cultures using CytoTox 96 nonradioactive cytotoxicity assay (Promega, Charbonnières-les-Bains, France) according to the manufacturer’s instructions. The absorbance of converted dye was measured at 490 nm with background subtraction at 690 nm.

### Statistical analysis

All statistical tests were done using the software Graph-Pad Prism 10.2.0.

## RESULTS

### Expression of NS1^CWA^ protein

Sequence comparison between NS1^CWA^ and NS1^MR766^ proteins identified seven amino acid differences (Table 1). These changes are distributed between the hydrophobic β-roll domain (amino-acids 1-29), the α*/*β Wing domain (amino-acids 38-151), the connector (amino-acids 152-180), and the β-ladder domain (amino-acids 181-352). We observed that NS1^CWA^ residues P92/K146/I162/R213 are unique as compared with NS1^MR766^ and NS1 proteins from viral strains of Asia/America lineage (Table 1).

**Table 1.** Amino-acid changes in ZIKV NS1 protein.

| NS1 |  | ZIKV of Africa lineage |  |  |  | ZIKV of Asia/America lineage |  |  |  |
| --- | --- | --- | --- | --- | --- | --- | --- | --- | --- |
| position | domain | ZIKV-15555<sup>1</sup> | SEN-2011<sup>2</sup> | SEN-2015<sup>3</sup> | MR766<sup>4</sup> | P6-740<sup>5</sup> | H/PF/2013<sup>6</sup> | BeH819015<sup>7</sup> | PVRABC59<sup>8</sup> |
| 21 | β-roll | V | V | V | I | V | V | V | V |
| 92 | $\alpha/\beta$ Wing | P | P | P | S | S | S | S | S |
| 146 | $\alpha/\beta$ Wing | K | K | K | E | E | E | E | E |
| 162 | connector | I | I | I | V | V | V | V | V |
| 191 | β-ladder | K | K | K | R | K | K | K | K |
| 213 | β-ladder | R | R | R | K | K | K | K | K |
| 286 | β-ladder | H | H | H | Y | H | H | H | H |

<sup>1</sup> GenBank access n°MN025403; <sup>2,3</sup> Viral strains Senegal-Kedougou 2011 (SEN-11) and Senegal-Kedougou 2015 (SEN-15) are referenced under ENA accession number n°PRJEB39677; <sup>4</sup> GenBank access n°LC002520; <sup>5</sup> GenBank access n°KX377336; <sup>6</sup> GenBank access n°KJ776791; <sup>7</sup> GenBank access n°KU365778; <sup>8</sup> GenBank access n°KX377337.

The PHYRE^2^ protein fold recognition server was used to test changes in ZIKV NS1 structure upon mutations (Fig. 1). Most of the amino acid changes between NS1^CWA^ and NS1^MR766^ are predicted to locate to disordered sequences. Only residues 213/286 are recruited into two distinct β-strands in β-ladder domain. The 3D structure prediction of ZIKV NS1 protein showed no apparent conformation changes between NS1^CWA^ and NS1^MR766^ (Fig.1).

**Figure 1.**
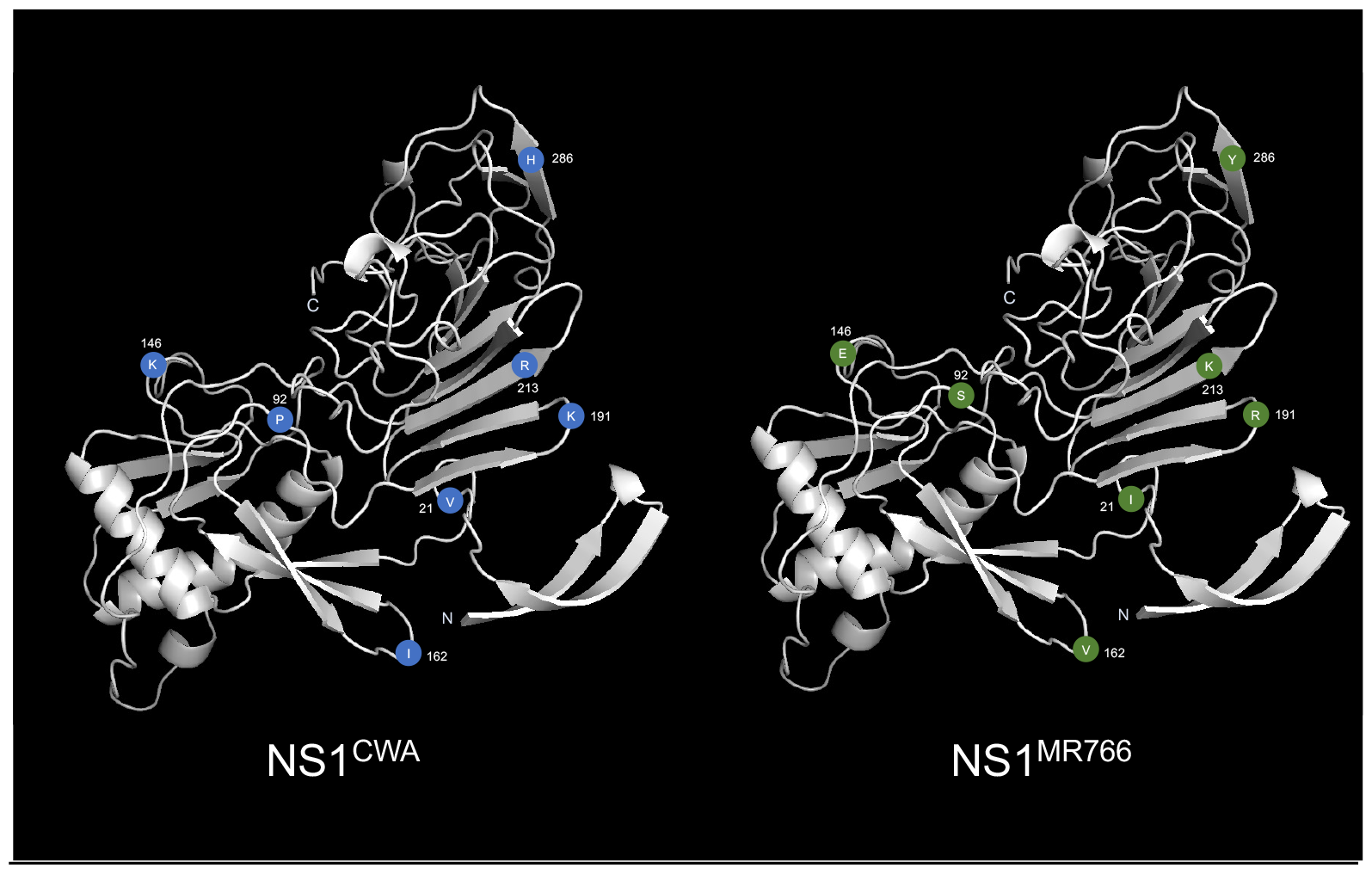
Three-dimensional structure prediction of contemporary West African ZIKV NS1 protein. The server Phyre^2^ was used to predict the 3D structure of NS1 proteins from Contemporary West Africa ZIKV (NS1^CWA^) and MR766 (NS1^MR766^). The prediction model is based on c5k6kb template (PDB: DOI: https://doi.org/10.2210/pdb5k6k/pdb) with 100% confidence and 97% coverage. The 3D viewing of the predicted structures was performed using PyMol molecular visualization system (version Z.1 INTEL-16.5.14). The positions of seven amino-acid residues that differentiate NS1^CWA^ from NS1^MR766^ are shown as colored closed circles.

We first examined whether NS1^CWA^ from NS1^MR766^ can impact protein expression. Synthetic genes coding for NS1 with optimized codons for expression in human cells were inserted into an expression vector (pcDNA3). In the resulting plasmids, the sequences coding for recombinant NS1 (rNS1) proteins were preceded by the authentic signal peptide corresponding to the second transmembrane domain of adjacent E protein and C-terminally tagged with a FLAG epitope. Given that the P92 residue might play a role in protein-protein interactions, site-directed mutagenesis was performed on plasmid pcDNA3/rNS1^MR766^ to generate a mutant plasmid bearing the S92P mutation. With respect to potential involvement of H286 in pH-dependent NS1 protein stability (23), the rNS1^MR766^ protein mutant also included the Y286H mutation.

Expression of rNS1 protein was analyzed in HEK-293T cells transfected for 48 h (Fig. 2A). FACS analysis using anti-FLAG antibody showed that the plasmids expressing NS1^CWA^, NS1^MR766^ or the NS1^MR766^ mutant bearing the (S92P, Y286H) mutations gave a similar percentage (nearly 50%) of HEK-293T cells positive for NS1 expression (Fig. 2A). The mean intensity fluorescence was comparable between the different rNS1 proteins. Thus, the ZIKV rNS1 expression constructs were suitable for further studies. The subcellular distribution of rNS1 proteins was examined in transfected HEK-293T cells by confocal immunofluorescence microscopy using anti-FLAG antibody (Fig. 2B). We observed that NS1^MR766^ protein mostly accumulated in the perinuclear region. In contrast, NS1^CWA^ protein and NS1^MR766^ mutant were seen to form multiple discrete foci in the cytoplasm These results suggest that NS1^CWA^ and NS1^MR766^ can vary in their subcellular distribution in HEK-293T cells. The residues P92 and H286 might play a role in the trafficking of NS1^CWA^ protein.

**Figure 2.**
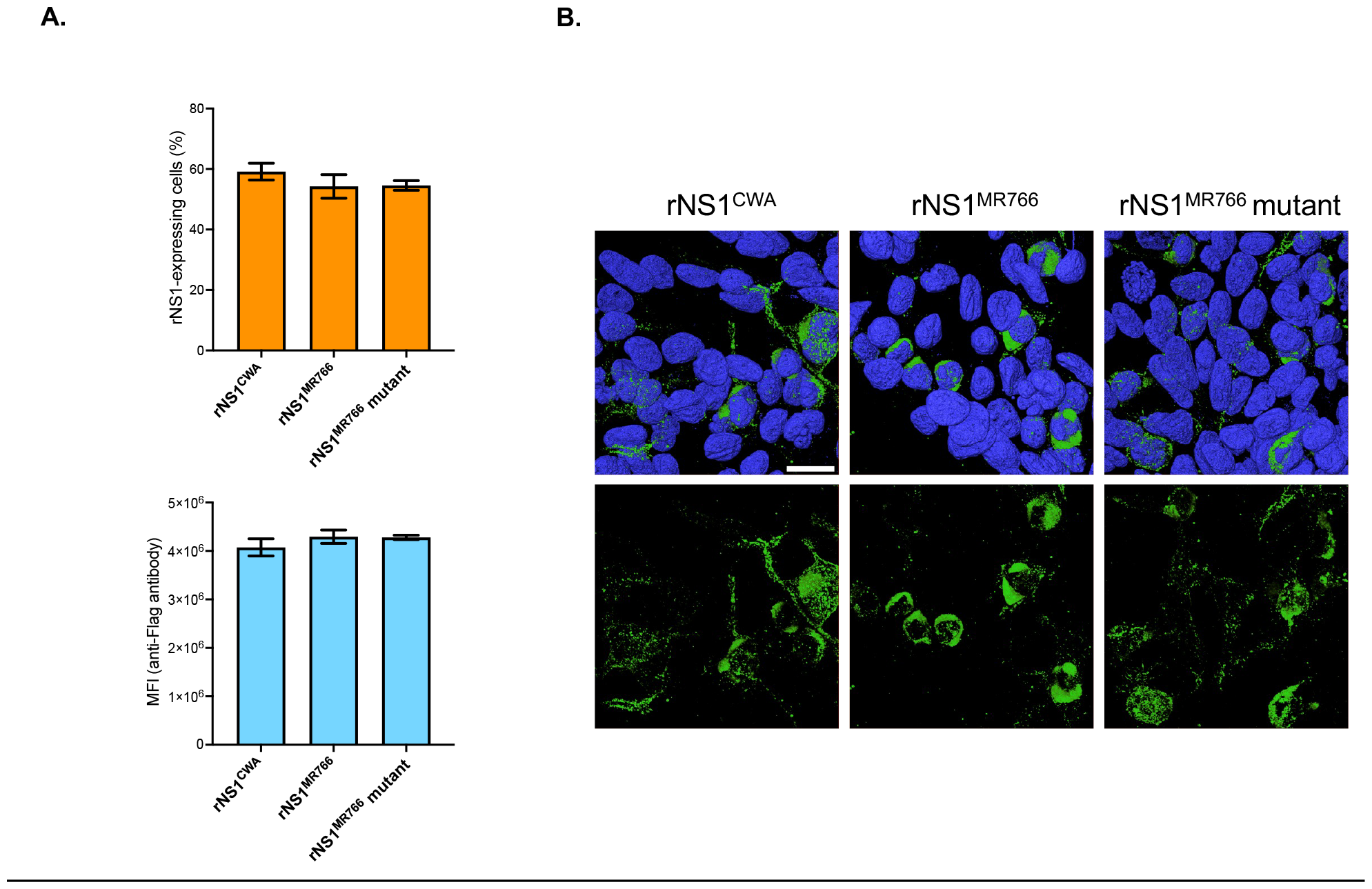
Immunodetection of rNS1 proteins. HEK-293T cells were transfected for 24h (B) or 48 h (A) with plasmids expressing rNS1^MR766^, rNS1^CWA^, or rNS1^MR766^-(P92, H286) mutant (rNS1^MR766^ mutant) or mock-transfected (control). *In* (**A**), Intracellular rNS1 proteins were labeled using anti-FLAG antibody. The percentage of cells positive for rNS1 expression and the mean fluorescence intensity (MFI) of FITC signal in positive cells were examine by FACS analysis. The results are the mean (± SEM) of three independent assays. No statistical differences were observed between the samples. In (**B**), three-dimensional visualization of rNS1 intracellular localization. HEK-293T cells were transfected for 24 h with plasmids expressing rNS1^MR766^, rNS1^CWA^, or rNS1^MR766^-(P92, H286) mutant (rNS1^MR766^ mutant). Cells were stained with anti-FLAG antibody (green) for confocal immunofluorescence analysis. Nuclei were strained with DAPI (blue). Cells surface rendering images are shown with (top) or without (bottom) their nucleus. The same magnification was used throughout. Scale bar, 25 µM.

Immunoblotting was performed on HEK-293T cell lysates expressing rNS1 proteins using anti-FLAG antibody and anti-*pan* flavivirus NS1 mAb 4G4 (Fig 3). Given that intracellular NS1 protein essentially exists as a heat-labile homo-dimer, cell lysates were first analyzed after heat denaturation (Fig. 3A). Anti-FLAG antibody detected comparable amounts of rNS1^CWA^, rNS1^MR766^, and rNS1^MR766^ mutant proteins which have similar migration profiles. As shown in Fig. 3B, anti-NS1 mAb 4G4 was able to detect dimeric forms of rNS1^CWA^, rNS1^MR766^, and rNS1^MR766^ mutant proteins (apparent MW estimated to 70 kDa in native samples). We noted that the antigenic reactivity of dimer and monomer rNS1^CWA^ proteins with conformation-specific mAb 4G4 was lower compared with rNS1^MR766^. The amino acid substitutions S92P and Y286H affected reactivity of rNS1^MR766^ with mAb 4G4 suggesting that the two residues may have an effect on the exposure of 4G4 antibody epitope. These results showed that mAb 4G4 is suitable to detect the soluble rNS1 released into the extracellular environment in HEK-293T cells. To estimate the secretion efficiency of rNS1 protein, cell supernatants were collected at 48 h post-transfection and then analyzed by a dot-blot assay using anti-NS1 4G4 mAb (Fig. 3C). A greater amount of extracellular NS1^CWA^ protein was detected in HEK-293T cell supernatant as compared with NS1^MR766^ or NS1^MR766^ mutant. By measuring signal intensity of dot-blot assays, we found that the amount of extracellular rNS1^CWA^ protein was at least 2-fold higher than that of rNS1^MR766^ (Fig. 3B,C). Immunoblot assay using anti-FLAG antibody verified that secretion rate of rNS1^CWA^ was significantly higher as compared with NS1^MR766^ (Fig. 3D). There was no effect of amino-acid substitutions S92P and Y286H on the secretion efficiency of rNS1^MR766^ protein (Fig. 3C, D).

**Figure 3.**
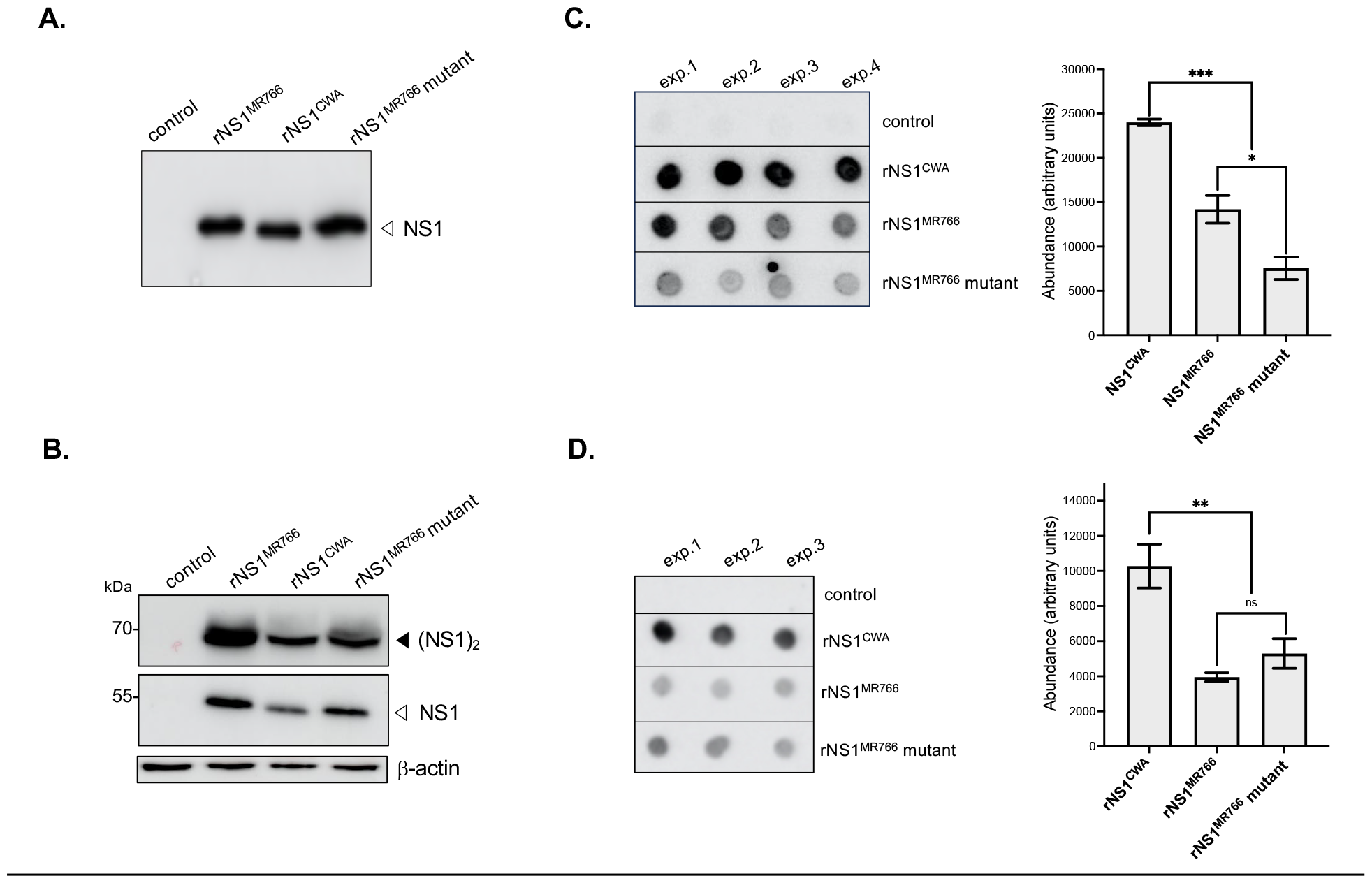
Expression of soluble rNS1 proteins. HEK-293T cells were transfected for 48 h with plasmids expressing rNS1^MR766^, rNS1^CWA^, or rNS1^MR766^-(P92, H286) mutant (rNS1^MR766^ mutant), or mock-transfected (control). *In* (**A**), immunoblot assay was performed on cell lysates by probing with anti-FLAG antibody. Samples were heated before loading on SDS-PAGE. The open head arrow indicates monomer NS1 protein. *In* (**B**), immunoblot assay was performed on cell lysates with anti-NS1 4G4 mAb. Samples were heated (bottom) or not (top) before loading on SDS-PAGE. The open head arrow indicates monomer NS1 protein. The β-actin was detected as protein-loading control for lysate samples. The close head arrow indicates dimer NS1 protein. In (**C**), cell supernatant samples from four independent transfection assays were analyzed by dot-blotting using anti-NS1 4G4 mAb. The signal intensity was quantified using Image J software to estimate the amounts of secreted soluble rNS1 protein. Statistical analysis of the mean (± SEM) of three replicates (** *P* < 0.01; ns: not significant). In (**D**), cell supernatant samples from three independent transfection assays were analyzed by dot-blotting using anti FLAG antibody. The signal intensity was quantified using Image J software to estimate the amounts of secreted soluble rNS1 protein. Statistical analysis of the mean (± SEM) of three replicates (** *P* < 0.01; ns: not significant).

To rule out that greater amount of extracellular rNS1^CWA^ protein reflects a higher cytotoxicity of the protein, lactate dehydrogenase (LDH) activity was measured in HEK-293T cells at 48 h post-transfection (Fig. 4). Although rNS1^CWA^ expression was slightly more cytotoxic than rNS1^MR766^, the effect of rNS1^CWA^ and rNS1^MR766^ mutant on cell viability was comparable despite the difference in protein secretion efficiency. Thus, it seems unlikely that secretion efficiency rNS1^CWA^ depended on increased cytotoxicity. Taken together, these results showed that rNS1^CWA^ and rNS1^MR766^ proteins differ in their subcellular distribution. The soluble rNS1^CWA^ protein has been found in a greater amount in the cell supernatant emphasizing a role for the residues that differentiate rNS1^CWA^ from rNS1^MR766^ in the secretion efficiency of the protein. The P92/H286 residues may play a role in subcellular distribution but not secretion efficiency of the rNS1^CWA^ protein.

**Figure 4.**
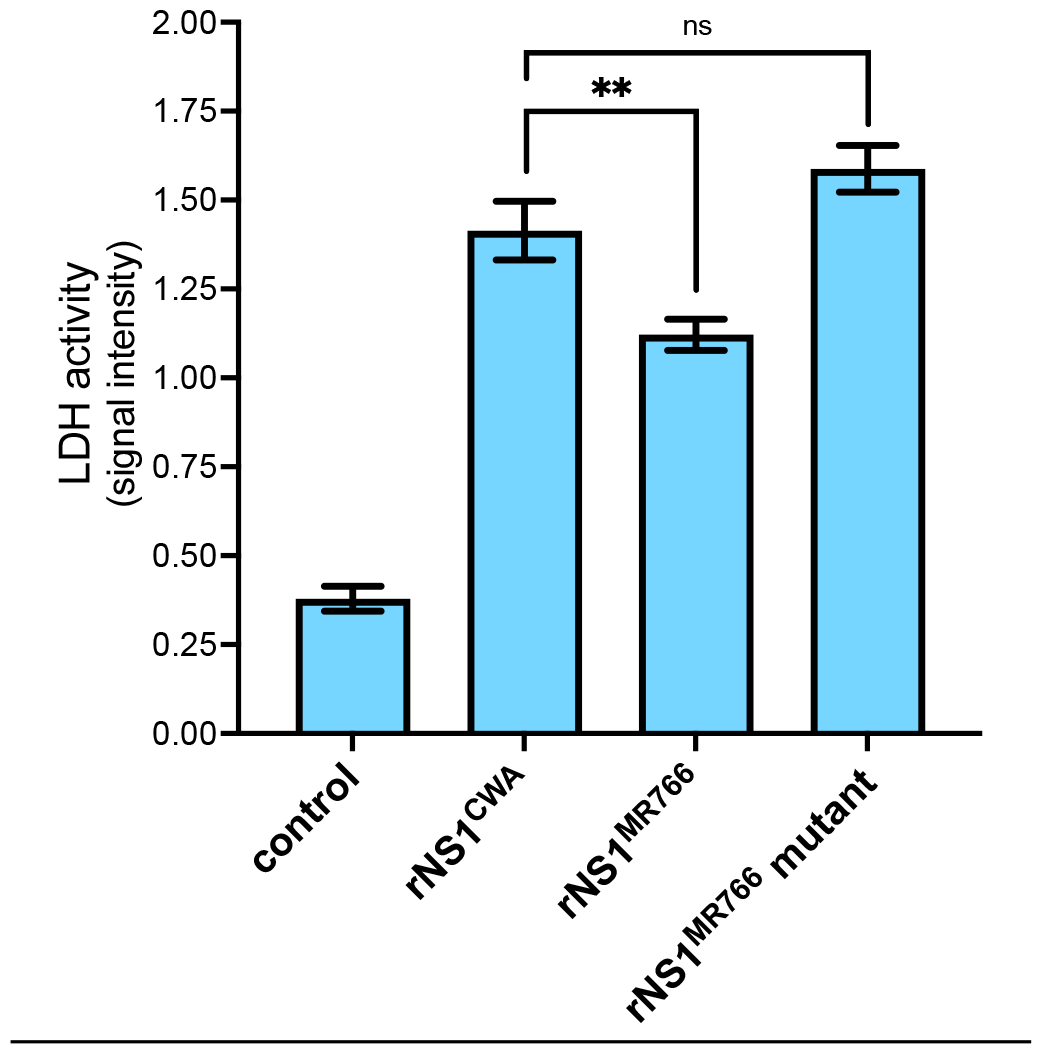
Cytotoxicity of rNS1 proteins. HEK-293T cells were transfected for 48 h with plasmids expressing rNS1^MR766^, rNS1^CWA^, or rNS1^MR766^-(P92, H286) mutant (rNS1^MR766^ mutant), or mock-transfected (control). LDH activity was measured and O.D. values were expressed as signal intensity. Statistical analysis of the mean (± SEM) of replicates (** *P* < 0.01; ns: not significant).

### Replication properties of a chimeric MR766/NS1^CWA^ virus

We next investigated whether the secretion efficiency of soluble NS1^CWA^ protein might have an effect on ZIKV replication. Consequently, the NS1^CWA^ gene from viral strain ZIKV-15555 (NS1^ZIKV-15555^) was introduced into available infectious molecular clone MR766^MC^ by reverse genetic using the ISA method (21). The resulting chimeric MR766^MC^/NS1^CWA^ virus includes the seven NS1 amino-acids substitutions that differentiate ZIKV-15555 from MR766^MC^. The chimeric MR766^MC^/NS1^ZIKV-15555^ virus was first assayed for virus replication in non-human primate Vero cells infected at m.o.i. of 0.1 (Fig. 5). The progeny production of MR766^MC^ chimera was increased by at least 10 fold at 48 h p.i. as compared with parental virus (Fig. 5A). This correlated with a rate of vRNA increased by 50% Fig. 5B) and higher percentage of ZIKV-infected cells (Fig. 5C). The measure of LDH release showed that infection with chimeric MR766^MC^/NS1^ZIKV-15555^ virus caused a more pronounced loss of cell viability compared to parental virus (Fig. 5D). Thus, chimeric MR766^MC^/NS1^CWA^ virus has a greater efficacy to replicate in non-human primate cells leading to an increased virus progeny production associated to more pronounced cytotoxicity.

**Figure 5.**
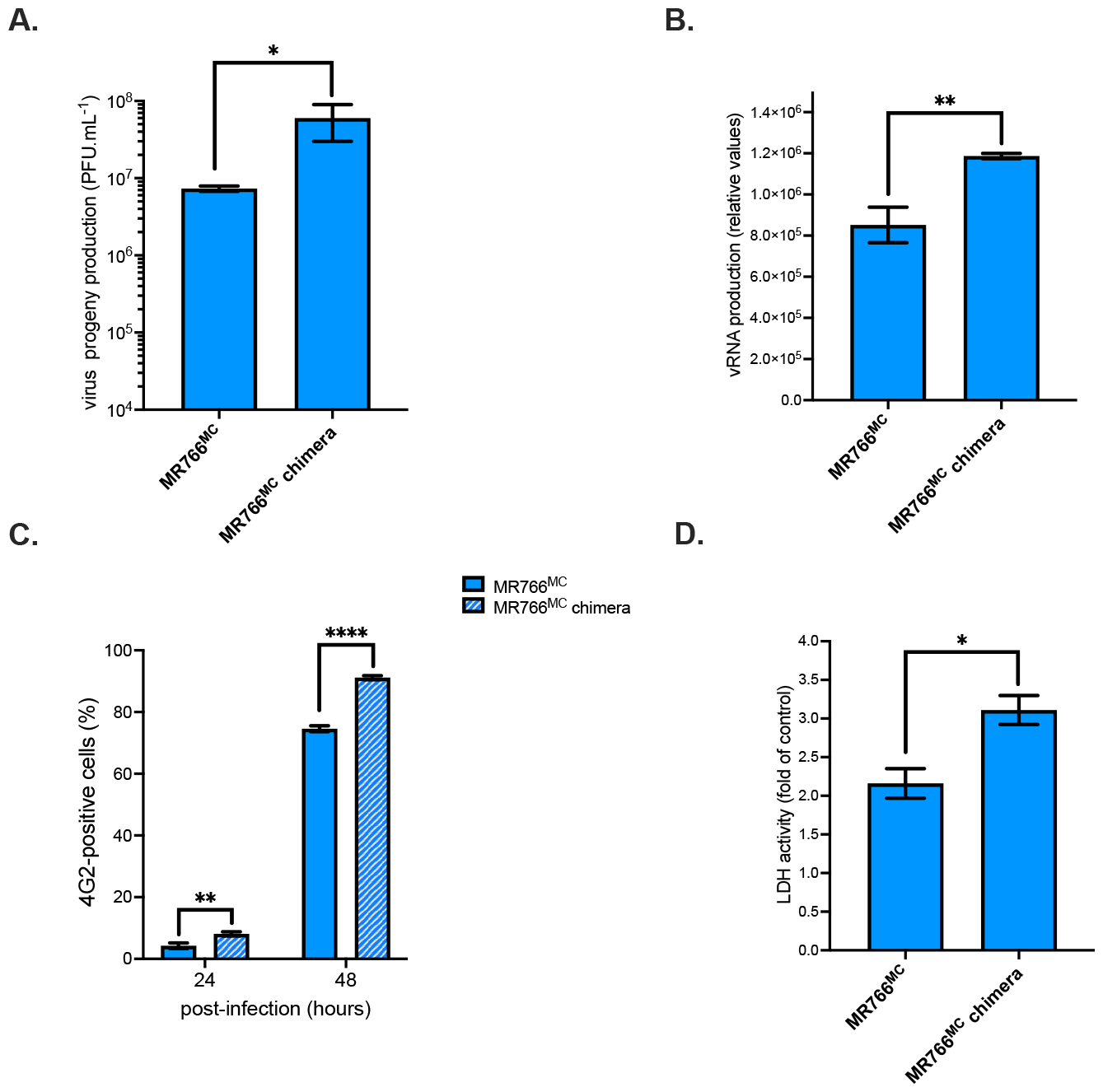
Replication of chimeric MR766^MC^/NS1^CWA^ virus in Vero cells. Vero cells were infected with MR766^MC^ or the chimeric MR766^MC^/NS1^CWA^ virus (MR766^MC^ chimera) at m.o.i. 0.1. In (**A**), virus progeny production (PFU.mL^-1^) was examined by plaque-forming assay. In (**B**), intracellular viral RNA production was determined by RT-qPCR at 48 h p.i. In (**C**), the percentage of ZIKV-infected cells based on FACS analysis using anti-E mAb 4G2. *In* (**D**), LDH activity were measured at 48h p.i and expressed as a percentage relative to mock-infected cells (control). Statistical analysis was noted (\*\*\**P* < 0.001, \*\**P* < 0.01, * *P* < 0.05).

The replication efficiency of chimeric MR766^MC^/NS1^CWA^ virus was next assessed in human epithelial A549 cells that have been chosen for their permissiveness to ZIKV infection (22,24) (Fig. 6). At 48 h p.i., there was an increase of nearly 1 log of progeny virus production for MR766^MC^ chimera compared to parental virus as it has been observed with non-human primate cells (Fig. 6A). The amount of vRNA in MR766^MC^ chimera-infected A549 cells was 10-fold higher (Fig. 6B) associated to a 2-fold increase in the percentage of infected cells as compared with MR766^MC^ (Fig. 6C). Infection with chimeric MR766^MC^/NS1^CWA^ virus resulted in a pronounced loss of cell viability at 48 h p.i. (Fig. 6D). Thus, MR766^MC^ chimera displays a greater growth capacity in A549 cells in comparison with parental virus. These results showed that insertion of NS1^ZIKV-15555^ protein in MR766^MC^ enhances viral replication associated to a more pronounced cell death. In conclusion, the seven residues that differentiate NS1^CWA^ from NS1^MR766^ enhance virus replication regardless of cell type.

**Figure 6.**
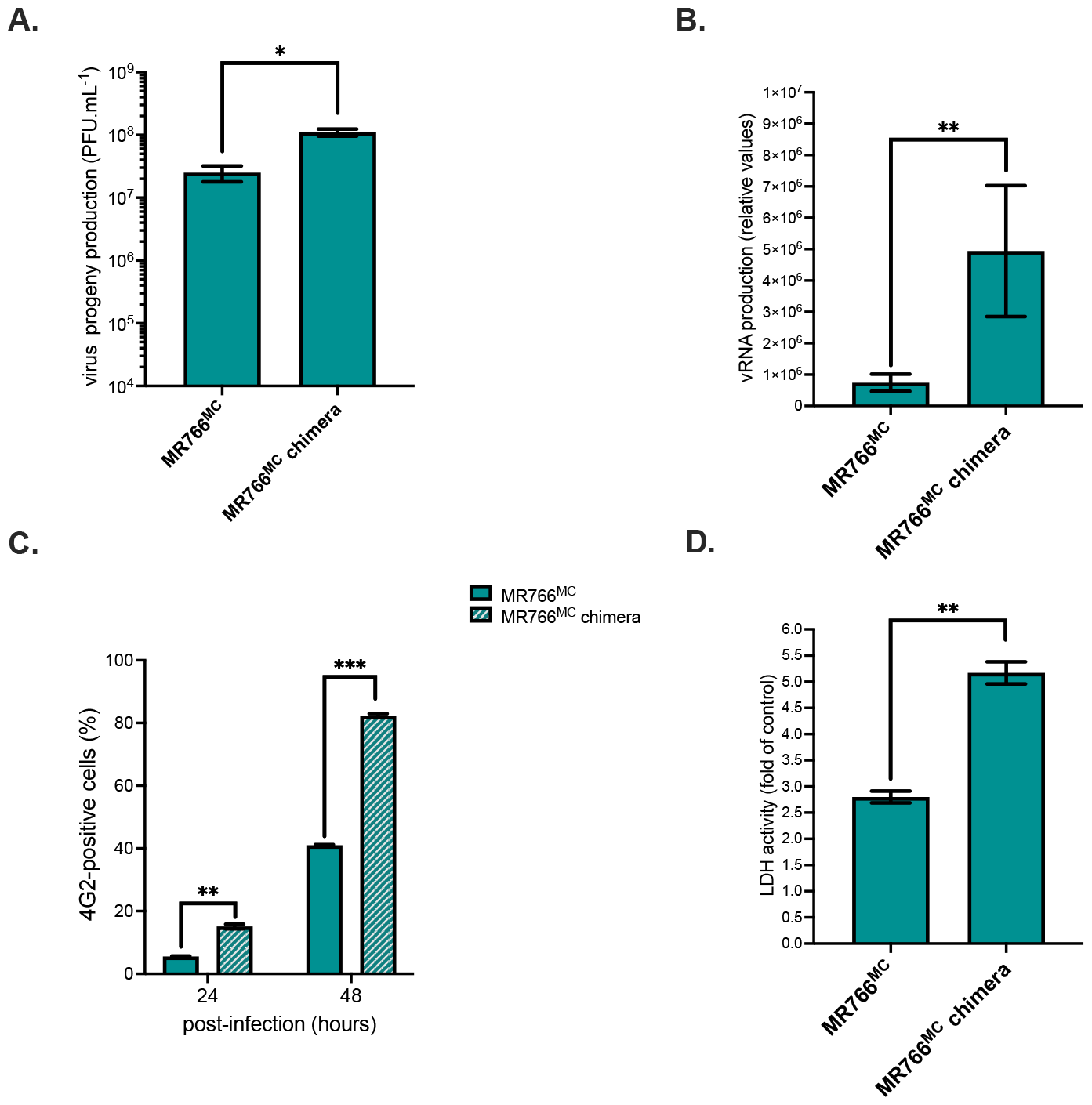
Replication of chimeric MR766^MC^/NS1^CWA^ virus in A549 cells. A549 cells were infected with MR766^MC^ or chimeric MR766^MC^/NS1^CWA^ virus (MR766^MC^ chimera) at m.o.i. 1. In (**A**), virus progeny production (PFU.mL^-1^) was examined using a conventional plaque-forming assay. In (**B**), intracellular viral RNA production was determined by RT-qPCR at 48 h p.i. *In* (**C**), the percentage of ZIKV-infected cells based on FACS analysis using anti-E mAb 4G2. In (**D**), LDH activity was measured at 48h p.i and expressed as a percentage relative to mock-infected cells (control). Statistical analysis was noted (\*\*\**P* < 0.001, \*\**P* < 0.01, * *P* < 0.05).

ZIKV infection results in activation of the intracellular signaling pathways leading to antiviral innate immune responses (25). We examined whether in the greater vulnerability of A549 cells to chimeric MR766^MC^/NS1^CWA^ virus also resulted in ISGs and IFN-β expression (Fig. 7). The relative abundance of IFN-β mRNA and various ISGs mRNAs was assessed by RT-qPCR on total RNA extracted from ZIKV-infected cells at 48 h p.i. (Fig. 7A). Infection with MR766^MC^ induced expression of *IFN-β* and ISGs with antiviral functions such as *Mx, OAS, IFIT, ISG15*, and *viperin*. Immunoblot assay with anti-IFIT1 or ISG15 antibody confirmed the upregulation of ISG expression during ZIKV infection (Fig. 7B). Importantly, we observed lower transcription levels of *IFN-β* and some ISGs in A549 cells infected by MR766^MC^ chimera compared to parental virus (Fig. 7A). Only *ISG15* and *OAS3* transcripts were not significantly different. The data indicated that NS1^CWA^ protein can down-regulate *IFN-β* and ISG induction in A549 cells infected by ZIKV.

**Figure 7.**
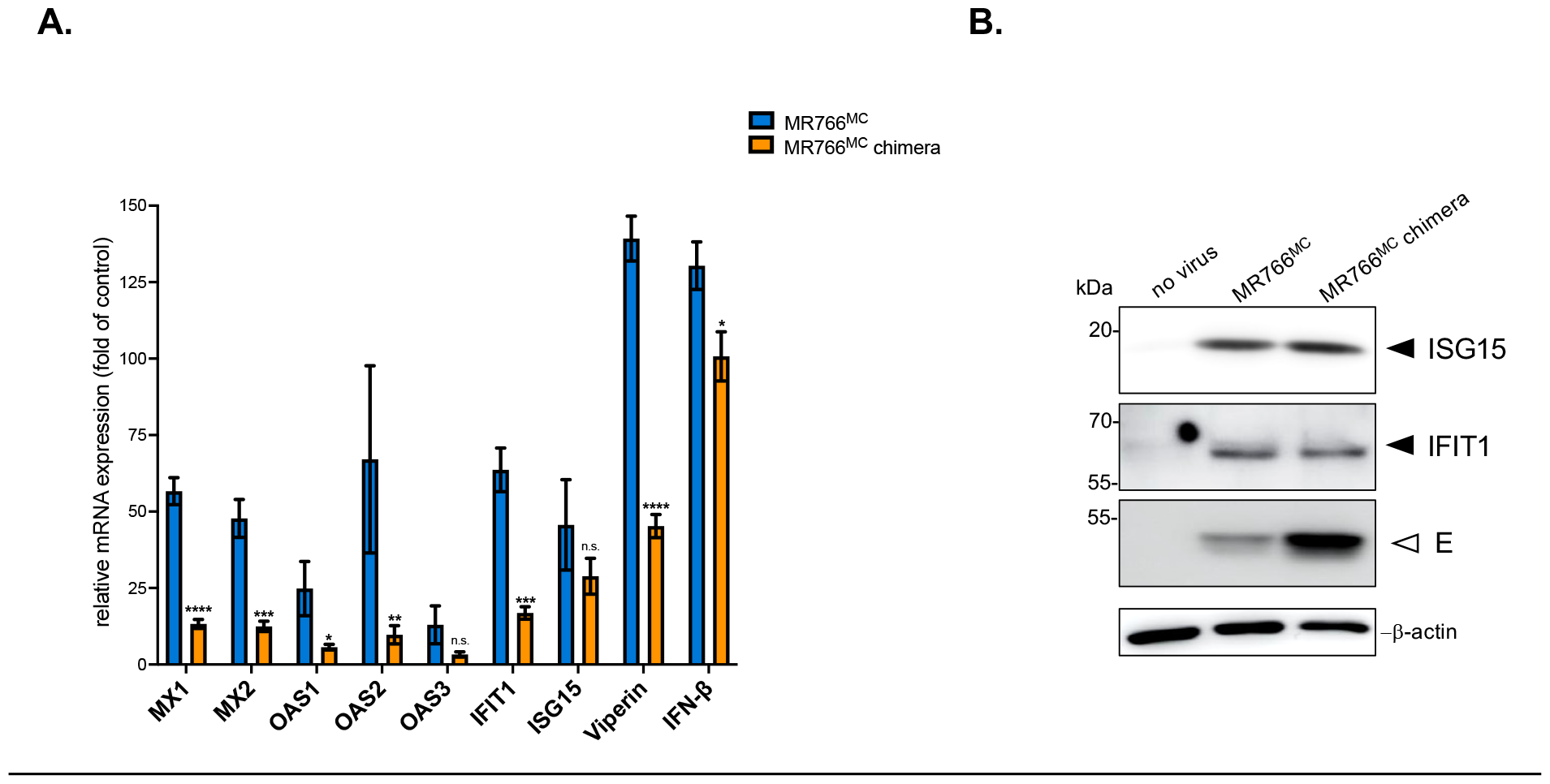
Induction of *IFN-β* and ISG expression in A549 cells infected by ZIKV. A549 cells were infected with MR766^MC^ or the chimeric MR766^MC^/NS1^CWA^ virus (MR766^MC^ chimera) at m.o.i. 1. In (**A**), the relative abundance of IFN-β and ISG mRNA was determined at 48 h p.i. by RT-qPCR. House-keeping RPLPO36B4 mRNA served as an internal reference. The results are the mean (± SD) of three replicates. Asterisks indicate that the differences between MR766^MC^ and MR766^MC^ chimera for each cellular factor are statistically significant, using the unpaired *t* test (\*\*\*\**P* < 0.0001, \*\*\**P* < 0.001, \*\**P* < 0.01, * *P* < 0.05). In (**B**), Immunoblot assay was performed on cell lysates using anti-ISG15 or anti-IFIT1 antibodies as indicated. Anti-E mAb 4G2 was used to detect ZIKV E protein. β-actin was detected as protein-loading control for lysate samples.

## CONCLUSION

The NS1 protein has been considered as a major viral target for inclusion in diagnosis, vaccine and antivirals development (26-30). A greater attention must be paid on recently isolated ZIKV of African lineage, with regards to their epidemic potential and capacity to be highly teratogenic in humans (6). To our knowledge, the present study is the first characterization of ZIKV NS1 protein from viral strains that have been recently isolated in West Africa. We find there is a remarkable conservation of NS1 amino-acid residues between ZIKV strains recently isolated in Senegal and Guinea. Among them, an infrequent proline has been identified at position 92 in the α*/*β Wing domain which shares structural homology with the retinoic acid-inducible gene I (RIG-I) and melanoma differentiation-associated protein 5 (MDA5) (31,32). Both RIG-I and MDA5 are double-stranded RNA sensors that engage signaling cascades leading to Type-I IFN genes activation (33). The NS1^CWA^ residue 92 contributes to an intertwined loop (amino-acids 90-130) containing hydrophobic residues that have been proposed to interact with host factors on plasma membrane (34). The presence of a proline at position NS1-92 enriched a potential proline-enriched sequence (amino-acid residues 92-112) that can recruit factors involved in regulation of signaling cascades (31).

The NS1^CWA^ protein differs from the historical African ZIKV strain MR766 (NS1^MR766^) by seven amino-acid substitutions. Analysis of recombinant ZIKV NS1 proteins expressed in HEK-293T cells revealed that NS1^CWA^ and NS1^MR766^ proteins can differ in their subcellular distribution. The NS1^CWA^ protein has been observed in discrete foci-like structures in the cytoplasm whereas NS1^MR766^ protein accumulated in the perinuclear region. This change in subcellular distribution was associated to a higher secretion efficiency of NS1^CWA^ protein compared with NS1^MR766^ protein. The amino-acid substitutions S92P and Y286H have been introduced in recombinant NS1^MR766^ protein and the protein mutant showed in subcellular distribution that resembles to NS1^CWA^ protein. The immune reactivity of mAb 4G4 was lower with NS1^CWA^ and at the lesser extent NS1^MR766^ protein mutant, as compared with ^MR766^ protein. The P92 and H286 residues may have an influence on the binding of this conformation-specific NS1 antibody even if the 3D prediction structure of ZIKV NS1 protein showed no obvious conformational changes between NS1^CWA^ and NS1^MR766^. Further studies of the impact of the conserved P92 residue on features of contemporary NS1 protein of West African ZIKV may help to clarify whether a proline at position 92 of the α*/*β wing domain alters NS1 subcellular distribution and antigenic reactivity of the protein. There was an increased secretion of NS1^CWA^ protein compared to NS1^MR766^ from human cells. Among the seven mutations that differentiate the two proteins, the residues P92 and H286 were not sufficient to influence the secretion efficiency of NS1^MR766^. It is possible that increased secretion of NS1^CWA^ protein was attributable to non-conservative amino-acid change E136K in the α*/*β wing domain. Introduction of NS1-E136K mutation in NS1^MR766^ protein could help interpret NS1^CWA^ residues in secretion efficiency of contemporary NS1 from West African ZIKV. In view of the importance of NS1 extracellular forms in the pathogenesis of ZIKV infection, understanding the consequences of an increased secretion of the protein is an issue that will be the subject of further investigation.

A chimeric virus derived from infectious molecular clone MR766^MC^ was engineered with the NS1^CWA^ gene from viral strain ZIKV-15555 in replacement of the authentic one. The viral growth of chimeric MR766^MC^/NS1^CWA^ virus was enhanced leading to an increased virus progeny production and higher cytotoxicity as compared with parental virus. The improved replication of chimeric MR766^MC^/NS1^CWA^ virus has been observed in Vero E6 and A549 cells. Given involvement of NS1 protein in viral RNA replication (11), NS1^CWA^ protein might have a greater propensity to interact with the other NS proteins thus enhancing activity of RCs (12). Also, NS1^CWA^ protein would be more efficient in the ER modeling and vesicle packet formation that are required for an effective viral RNA replication (35,36). We cannot rule out that increased secretion efficacy of NS1^CWA^ protein makes viral morphogenesis easier (36). We showed that MR766^MC^ infection induces *IFN-β* expression in A549 cells. There was a lower magnitude of *IFN-β* mRNA up-regulation in A549 cells infected by chimeric MR766^MC^/NS1^CWA^ virus. Whether the efficiency of NS1^CWA^ protein to antagonize Type-I IFN induction contributes to efficient virus replication remains to be elucidated. IFN-β Induces the expression of a large numbers of ISGs through a Janus kinase/signal transducer and activator of transcription (JAK/STAT) pathway (37). Among ISGs with antiviral functions, we found that MR766^MC^ induced *MX, OAS, IFIT1, viperin* and *ISG15* mRNA expression in A549 cells. It has been reported that IFIT1 interacts with STING/MITA to negatively regulates IRF3 activation and also inhibits the mRNA translation to restrain orthoflavivirus replication (38). Viperin has been identified as a one of the major ISG in control of ZIKV infection through inhibition of viral RNA translation (39-42). Infection with MR766^MC^ chimera expressing NS1^CWA^ protein resulted to a lower magnitude of *IFIT1* and *Viperin* mRNA induction in A549 cells compared to parental virus. Insertion of NS1^CWA^ protein in MR766^MC^ caused no change in ISG15 expression consisting with the notion that ZIKV exploits ISG15 as a negative regulator of IFN-type I signaling to benefit its replication (37). Whether NS1^CWA^ protein manipulates some ISGs with antiviral functions by targeting JAK/STAT-mediated downstream events that are essential for activation of the IFN-β responsive genes are a critical issue that remains to be investigated. Further studies should be carried out to develop an understanding of mechanisms by which NS1 protein of recently isolated African ZIKV strains manipulates innate immunity to promote viral replication. The data would expand our current knowledge on the pathogenic properties of contemporary ZIKV strains from West Africa (6,21).

## Acknowledgements

We do want to thank L. Lambrechts for his input and insight into sequence data for ZIKV strains Senegal-Kedougou. We thank J. Andries and N. Masri for technical assistance and PIMIT members for helpful discussions. We thank PICT platform (University of Reims Champagne-Ardenne) for imaging core facilities.

## Author contributions

D.M., P.D. and M.R conceived and designed the experiments. D.M., E.O. M-P.C and M.R performed the experiments. D.M., M-P.C, P.D., E.O., A.K and M.R analyzed the data. P.D. contributed to reagents/materials/analysis tools. D.M., P.D., and M.R wrote the paper. All authors have read and agreed to the published version of the manuscript.

---

**Table S1.**
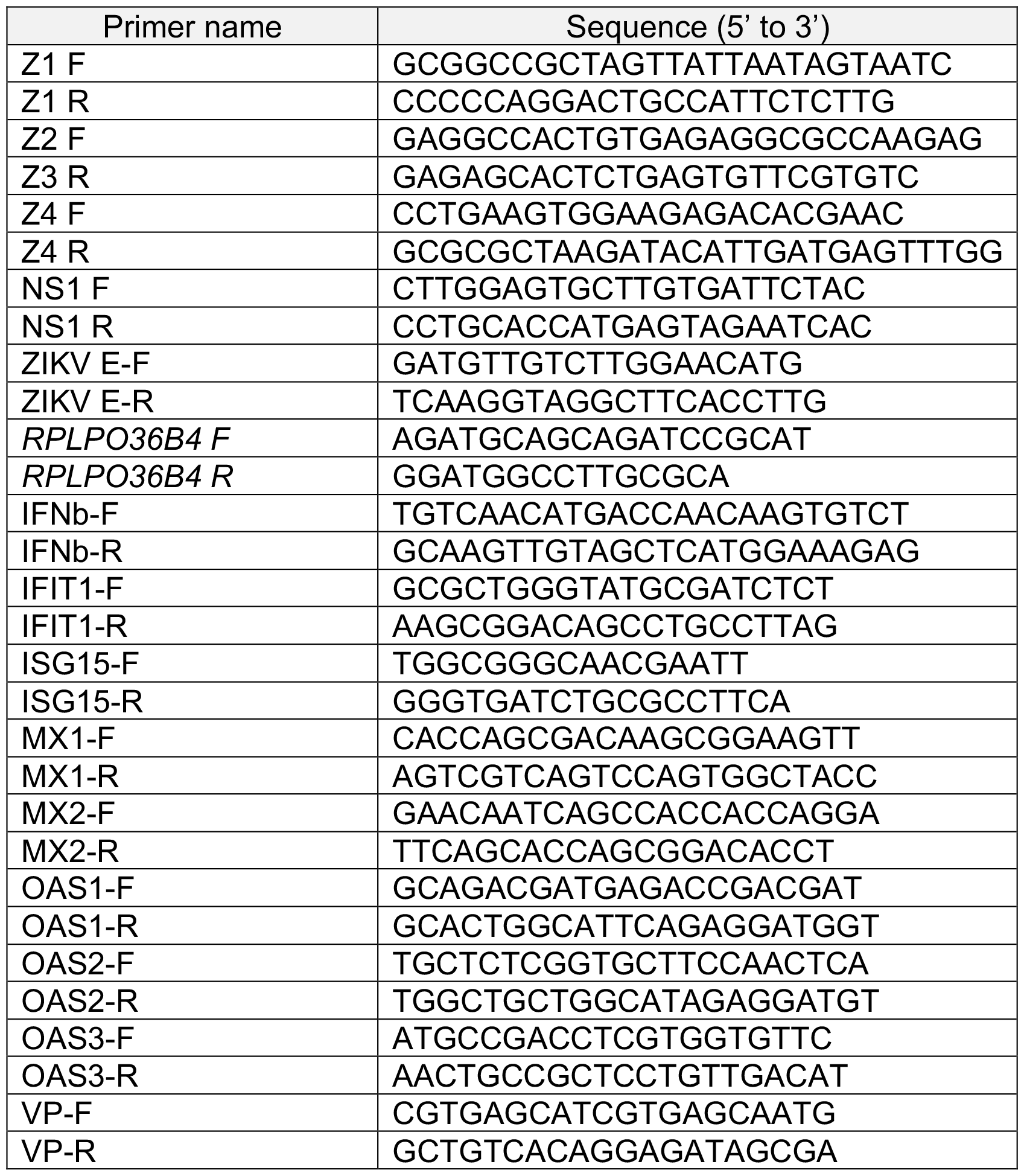
Sequences of primers for ISA method and RT-qPCR used in this study.

## REFERENCES

1. Pierson TC, Diamond MS. 2018. The emergence of Zika virus and its new clinical syndromes. Nature 560: 573–581.

2. Pierson TC, Diamond MS. 2020. The continued threat of emerging flaviviruses. NatMicrobiol 5: 796–812.

3. Beaver JT, Lelutiu N, Habib R, Skountzou I. 2018. Evolution of Two Major Zika Virus Lineages: Implications for Pathology, Immune Response, and Vaccine Development. Front Immunol 9:1640.

4. Freitas DA, Souza-Santos R, Carvalho LMA, Barros WB, Neves LM, Brasil P, Wakimoto MD. 2020. Congenital Zika syndrome: A systematic review. PLoS ONE 15:e0242367.

5. Vhp L, Aragão MM, Pinho RS, Hazin AN, Paciorkowski AR, Penalva de Oliveira AC, Masruha MR. 2020. Congenital Zika Virus Infection: a Review with Emphasis on the Spectrum of Brain Abnormalities. Curr Neurol Neurosci Rep 20:49.

6. Aubry F, Jacobs S, Darmuzey M, Lequime S, Delang L, Fontaine A, Jupatanakul N, Miot EF, Dabo S, Manet C, Montagutelli X, Baidaliuk A, Gámbaro F, Simon-Lorière E, Gilsoul M, Romero-Vivas CM, Cao-Lormeau V-M, Jarman RG, Diagne CT, Faye O, Faye O, Sall AA, Neyts J, Nguyen L, Kaptein SJF, Lambrechts L. 2021. Recent African strains of Zika virus display higher transmissibility and fetal pathogenicity than Asian strains. Nat Commun 12:916.

7. Rosinski JR, Raasch LE, Barros Tiburcio P, Breitbach ME, Shepherd PM, Yamamoto K, Razo E, Krabbe NP, Bliss MI, Richardson AD, Einwalter MA, Weiler AM, Sneed EL, Fuchs KB, Zeng X, Noguchi KK, Morgan TK, Alberts AJ, Antony KM, Kabakov S, Ausderau KK, Bohm EK, Pritchard JC, Spanton RV, Ver Hoove JN, Kim CBY, Nork TM, Katz AW, Rasmussen CA, Hartman A, Mejia A, Basu P, Simmons HA, Eickhoff JC, Friedrich TC, Aliota MT, Mohr EL, Dudley DM, O’Connor DH, Newman CM. 2023. Frequent first-trimester pregnancy loss in rhesus macaques infected with African-lineage Zika virus. PLoS Pathog 19:e1011282.

8. Aubry F, Dabo S, Manet C, Filipović I, Rose NH, Miot EF, Martynow D, Baidaliuk A, Merkling SH, Dickson LB, Crist AB, Anyango VO, Romero-Vivas CM, Vega-Rúa A, Dusfour I, Jiolle D, Paupy C, Mayanja MN, Lutwama JJ, Kohl A, Duong V, Ponlawat A, Sylla M, Akorli J, Otoo S, Lutomiah J, Sang R, Mutebi J-P, Cao-Lormeau V-M, Jarman RG, Diagne CT, Faye O, Faye O, Sall AA, McBride CS, Montagutelli X, Rašić G, Lambrechts L. 2020. Enhanced Zika virus susceptibility of globally invasive Aedes aegypti populations. Science 370:991–996.

9. Gomard Y, Lebon C, Mavingui P, Atyame CM. 2020. Contrasted transmission efficiency of Zika virus strains by mosquito species Aedes aegypti, Aedes albopictus and Culex quinquefasciatus from Reunion Island. Parasites Vectors 13:398.

10. Golubeva VA, Nepomuceno TC, Gregoriis G de, Mesquita RD, Li X, Dash S, Garcez PP, Suarez-Kurtz G, Izumi V, Koomen J, Carvalho MA, Monteiro ANA. 2020. Network of Interactions between ZIKA Virus Non-Structural Proteins and Human Host Proteins. Cells 9:153.

11. Akey DL, Brown WC, Jose J, Kuhn RJ, Smith JL. 2015. Structure-guided insights on the role of NS1 in flavivirus infection. Bioessays 37:489–494.

12. Mackenzie JM, Jones MK, Young PR. 1996. Immunolocalization of the dengue virus nonstructural glycoprotein NS1 suggests a role in viral RNA replication. Virology 220:232–240.

13. Shu B, Ooi JSG, Tan AWK, Ng T-S, Dejnirattisai W, Mongkolsapaya J, Fibriansah G, Shi J, Kostyuchenko VA, Screaton GR, Lok S-M. 2022. CryoEM structures of the multimeric secreted NS1, a major factor for dengue hemorrhagic fever. Nat Commun 13:6756.

14. Noisakran S, Dechtawewat T, Rinkaewkan P, Puttikhunt C, Kanjanahaluethai A, Kasinrerk W, Sittisombut N, Malasit P. 2007. Characterization of dengue virus NS1 stably expressed in 293T cell lines. J Virol Methods 142:67–80.

15. Smith JL. 2022. Flavivirus NS1: Structure and Function of an Enigmatic Virulence Factor. The FASEB Journal 36:fasebj.2022.36.S1.0I225.

16. Bos S, Poirier-Beaudouin B, Seffer V, Manich M, Mardi C, Desprès P, Gadea G, Gougeon M-L. 2020. Zika Virus Inhibits IFN-α Response by Human Plasmacytoid Dendritic Cells and Induces NS1-Dependent Triggering of CD303 (BDCA-2) Signaling. Front Immunol 11:582061.

17. Xia H, Luo H, Shan C, Muruato AE, Nunes BTD, Medeiros DBA, Zou J, Xie X, Giraldo MI, Vasconcelos PFC, Weaver SC, Wang T, Rajsbaum R, Shi P-Y. 2018. An evolutionary NS1 mutation enhances Zika virus evasion of host interferon induction. Nat Commun 9:414.

18. Shi Y, Dai L, Song H, Gao GF. 2018. Structures of Zika Virus E & NS1: Relations with Virus Infection and Host Immune Responses. Adv Exp Med Biol 1062:77–87.

19. Wilson JR, de Sessions PF, Leon MA, Scholle F. 2008. West Nile virus nonstructural protein 1 inhibits TLR3 signal transduction. J Virol 82:8262–8271.

20. Liu Y, Liu J, Du S, Shan C, Nie K, Zhang R, Li X-F, Zhang R, Wang T, Qin C-F, Wang P, Shi P-Y, Cheng G. 2017. Evolutionary enhancement of Zika virus infectivity in Aedes aegypti mosquitoes. Nature 545:482–486.

21. Gadea G, Bos S, Krejbich-Trotot P, Clain E, Viranaicken W, El-Kalamouni C, Mavingui P, Desprès P. 2016. A robust method for the rapid generation of recombinant Zika virus expressing the GFP reporter gene. Virology 497:157–162.

22. Machmouchi D, Courageot M-P, El-Kalamouni C, Kohl A, Desprès P. 2024. The replication properties of a contemporary Zika virus from West Africa depends on NS1/NS4B proteins 10.1101/2024.03.14.584947.

23. Poveda-Cuevas SA, Etchebest C, Barroso da Silva FL. 2018. Insights into the ZIKV NS1 Virology from Different Strains through a Fine Analysis of Physicochemical Properties. ACS Omega 3:16212–16229.

24. Frumence E, Roche M, Krejbich-Trotot P, El-Kalamouni C, Nativel B, Rondeau P, Missé D, Gadea G, Viranaicken W, Desprès P. 2016. The South Pacific epidemic strain of Zika virus replicates efficiently in human epithelial A549 cells leading to IFN-β production and apoptosis induction. Virology 493:217–226.

25. Hu H, Feng Y, He M-L. 2023. Targeting Type I Interferon Induction and Signaling: How Zika Virus Escapes from Host Innate Immunity. Int J Biol Sci 19:3015–3028.

26. Modhiran N, Song H, Liu L, Bletchly C, Brillault L, Amarilla AA, Xu X, Qi J, Chai Y, Cheung STM, Traves R, Setoh YX, Bibby S, Scott CAP, Freney ME, Newton ND, Khromykh AA, Chappell KJ, Muller DA, Stacey KJ, Landsberg MJ, Shi Y, Gao GF, Young PR, Watterson D. 2021. A broadly protective antibody that targets the flavivirus NS1 protein. Science 371:190–194.

27. Grubor-Bauk B, Wijesundara DK, Masavuli M, Abbink P, Peterson RL, Prow NA, Larocca RA, Mekonnen ZA, Shrestha A, Eyre NS, Beard MR, Gummow J, Carr J, Robertson SA, Hayball JD, Barouch DH, Gowans EJ. 2019. NS1 DNA vaccination protects against Zika infection through T cell-mediated immunity in immunocompetent mice. Sci Adv 5:eaax2388.

28. Brault AC, Domi A, McDonald EM, Talmi-Frank D, McCurley N, Basu R, Robinson HL, Hellerstein M, Duggal NK, Bowen RA, Guirakhoo F. 2017. A Zika Vaccine Targeting NS1 Protein Protects Immunocompetent Adult Mice in a Lethal Challenge Model. Sci Rep 7:14769.

29. Carpio KL, Barrett ADT. 2021. Flavivirus NS1 and Its Potential in Vaccine Development. Vaccines (Basel) 9:622.

30. Morales SV, Coelho GM, Ricciardi-Jorge T, Dorl GG, Zanluca C, Duarte Dos Santos CN. 2024. Development of a quantitative NS1 antigen enzyme-linked immunosorbent assay (ELISA) for Zika virus detection using a novel virus-specific mAb. Sci Rep 14:2544.

31. Kay BK, Williamson MP, Sudol M. 2000. The importance of being proline: the interaction of proline-rich motifs in signaling proteins with their cognate domains. FASEB J 14:231–241.

32. Lo NTN, Roodsari SZ, Tin NL, Wong MP, Biering SB, Harris E. 2022. Molecular Determinants of Tissue Specificity of Flavivirus Nonstructural Protein 1 Interaction with Endothelial Cells. J Virol 96:e0066122.

33. Rehwinkel J, Gack MU. 2020. RIG-I-like receptors: their regulation and roles in RNA sensing. Nat Rev Immunol 20:537–551.

34. Xu X, Song H, Qi J, Liu Y, Wang H, Su C, Shi Y, Gao GF. 2016. Contribution of intertwined loop to membrane association revealed by Zika virus full-length NS1 structure. EMBO J 35:2170–2178.

35. Eyre NS, Johnson SM, Eltahla AA, Aloi M, Aloia AL, McDevitt CA, Bull RA, Beard MR. 2017. Genome-Wide Mutagenesis of Dengue Virus Reveals Plasticity of the NS1 Protein and Enables Generation of Infectious Tagged Reporter Viruses. J Virol 91:e01455–17.

36. Tamura T, Torii S, Kajiwara K, Anzai I, Fujioka Y, Noda K, Taguwa S, Morioka Y, Suzuki R, Fauzyah Y, Ono C, Ohba Y, Okada M, Fukuhara T, Matsuura Y. 2022. Secretory glycoprotein NS1 plays a crucial role in the particle formation of flaviviruses. PLoS Pathog 18:e1010593.

37. Wang Y, Ren K, Li S, Yang C, Chen L. 2020. Interferon stimulated gene 15 promotes Zika virus replication through regulating Jak/STAT and ISGylation pathways. Virus Res 287:198087.

38. Fensterl V, Sen GC. 2015. Interferon-induced Ifit proteins: their role in viral pathogenesis. J Virol 89:2462–2468.

39. Van Der Hoek KH, Eyre NS, Shue B, Khantisitthiporn O, Glab-Ampi K, Carr JM, Gartner MJ, Jolly LA, Thomas PQ, Adikusuma F, Jankovic-Karasoulos T, Roberts CT, Helbig KJ, Beard MR. 2017. Viperin is an important host restriction factor in control of Zika virus infection. Sci Rep 7:4475.

40. Vanwalscappel B, Gadea G, Desprès P. 2019. A Viperin Mutant Bearing the K358R Substitution Lost its Anti-ZIKA Virus Activity. Int J Mol Sci 20:1574.

41. Rivera-Serrano EE, Gizzi AS, Arnold JJ, Grove TL, Almo SC, Cameron CE. 2020. Viperin Reveals Its True Function. Annu Rev Virol 7:421–446.

42. Hsu JC-C, Laurent-Rolle M, Pawlak JB, Xia H, Kunte A, Hee JS, Lim J, Harris LD, Wood JM, Evans GB, Shi P-Y, Grove TL, Almo SC, Cresswell P. 2022. Viperin triggers ribosome collision-dependent translation inhibition to restrict viral replication. Mol Cell 82:1631-1642.e6.

